# Heritability Estimation and Differential Analysis with Generalized Linear Mixed Models in Genomic Sequencing Studies

**DOI:** 10.1101/359265

**Authors:** Shiquan Sun, Jiaqiang Zhu, Sahar Mozaffari, Carole Ober, Mengjie Chen, Xiang Zhou

## Abstract

**Motivation:** Genomic sequencing studies, including RNA sequencing and bisulfite sequencing studies, are becoming increasingly common and increasingly large. Large genomic sequencing studies open doors for accurate molecular trait heritability estimation and powerful differential analysis. Heritability estimation and differential analysis in sequencing studies requires the development of statistical methods that can properly account for the count nature of the sequencing data and that are computationally efficient for large data sets.

**Results:** Here, we develop such a method, PQLseq (Penalized Quasi-Likelihood for sequencing count data), to enable effective and efficient heritability estimation and differential analysis using the generalized linear mixed model framework. With extensive simulations and comparisons to previous methods, we show that PQLseq is the only method currently available that can produce unbiased heritability estimates for sequencing count data. In addition, we show that PQLseq is well suited for differential analysis in large sequencing studies, providing calibrated type I error control and more power compared to the standard linear mixed model methods. Finally, we apply PQLseq to perform gene expression heritability estimation and differential expression analysis in a large RNA sequencing study in the Hutterites.

**Availability and implementation:** PQLseq is implemented as an R package with source code freely available at www.xzlab.org/software.html and https://cran.r-project.org/web/packages/PQLseq/index.html.

**Contact:** XZ

**Supplementary information:** Supplementary data are available online.

## INTRODUCTION

Generalized linear mixed model (GLMM) has recently emerged as a powerful statistical tool for the analysis of high throughput genomics sequencing studies (Lea, et al., 2015; Sun, et al., 2017; Weissbrod, et al., 2017; Zhang, et al., 2017). The main application of GLMM in these genomic sequencing studies is so far restricted to differential analysis, which aims to identify genomic units (e.g. genes or CpG sites) that are associated with a predictor of interest (e.g. disease status, treatment, environmental covariates, or genotypes). Common analysis examples include differential expression analysis in RNA sequencing (RNAseq) studies (Conesa, et al., 2016; Pickrell, et al., 2010) and differential methylation analysis in bisulfite sequencing (BSseq) studies (Irizarry, et al., 2009; Oakes, et al., 2016). Effective differential analysis with sequencing data often requires statistical methods to both account for the count nature of sequencing data and effectively control for sample non-independence – a common phenomenon in sequencing studies caused by individual relatedness, population structure, or hidden confounding factors (Dubin, et al., 2015; Scott, et al., 2016; Tung, et al., 2015). GLMM accomplishes both tasks by relying on exponential family distributions to directly model sequencing count data and by introducing random effects terms to account for sample non-independence. In effect, GLMM generalizes both the linear mixed model (LMM) that has been widely used to control for sample non-independence in association studies (Kang, et al., 2010; Lippert, et al., 2011; Zhou and Stephens, 2012), and over-dispersed count models (e.g. negative-binomial, beta-binomial) that have been widely used for differential analysis in sequencing studies (Love, et al., 2014; Robinson, et al., 2010; Sun, et al., 2014). By combining the benefits of the two commonly used methods, GLMM properly controls type I error and improves power for differential analysis (Lea, et al., 2015; Sun, et al., 2017).

While the existing applications of GLMM in genomic sequencing studies have been primarily restricted to differential analysis, the similarity between GLMM and LMM begs the question on whether GLMM can also be applied to estimate heritability for sequencing count data. Heritability measures the proportion of phenotypic variance explained by genetics and is an important quantity that facilitates the understanding the genetic basis of phenotypic variation. The standard tool for estimating heritability is LMM, which has long been applied for heritability estimation (Abecasis, et al., 2000; Almasy and Blangero, 1998; Amos, 1994; Diao and Lin, 2006; Visscher, et al., 2008; Zhou, 2017) or SNP heritability estimation (de los Campos, et al., 2015; Wray, et al., 2013; Yang, et al., 2010; Zhou, 2017; Zhou, et al., 2013) for various quantitative traits in the setting of genome-wide association studies (GWASs). In the setting of genomics studies, LMM has also been recently applied to estimate gene expression heritability (Emilsson, et al., 2008; Monks, et al., 2004; Price, et al., 2011; Tung, et al., 2015; Wright, et al., 2014), methylation level heritability (Banovich, et al., 2014; Bell, et al., 2012; McRae, et al., 2014), as well as various other molecular traits heritability (Cheng, et al., 2017). However, LMM is specifically designed for analyzing quantitative traits. In genomic sequencing studies, the application of LMM requires *a priori* transformation of the count data to continuous data before heritability estimation (Tung, et al., 2015; Wheeler, et al., 2016). Transforming sequencing count data may fail to properly account for the sampling noise from the underlying count generating process, and may inappropriately attribute such noise to independent environmental variation -- thus running the risk of overestimating environmental variance and subsequently underestimating heritability. In contrast, GLMM directly models count data, and as will be shown in the present study, has the potential to become a more accurate alternative than LMM for heritability estimation in genomic sequencing studies.

Both the above two applications of GLMM for differential analysis and heritability estimation require accurate and scalable inference algorithms to accommodate the increasingly large genomic sequencing studies that are being collected today. Indeed, several genomic projects have already collected sequencing data on hundreds of individuals (Ardlie, et al., 2015; Battle, et al., 2014; Consortium, 2015), and the recent TOPMed omics sequencing project further aims to sequence a few thousands of individuals in the next couple of years. Compared with small sample studies, large genomic sequencing studies are better powered and more reproducible, and are thus becoming increasingly common in genomics. In addition, large-scale population sequencing studies pave ways for accurate estimation of heritability for various molecular traits. Unfortunately, existing algorithms for fitting GLMM in genomic sequencing studies are not scalable. In addition, as will be shown in the present study, existing GLMM algorithms do not always produce calibrated *p*-values for differential analysis nor accurate heritability estimates.

In terms of scalability, existing algorithms to fit GLMM are generally computationally expensive due to an intractable high-dimensional integral in the GLMM likelihood (Breslow and Clayton, 1993; Chen, et al., 2016; Weissbrod, et al., 2017). For example, the frequentist method Mixed model Association for Count data via data AUgmentation algorithm (MACAU) relies on a Bayesian strategy of Markov chain Monte Carlo (MCMC) sampling to numerically approximate the integration in GLMM. However, though accurate, MCMC based strategy is computationally inefficient for large sample size: it takes MACAU several days to analyze moderate-sized RNAseq or BSseq data with a few hundred individuals. To overcome the computational bottleneck of MCMC-based approaches, recent studies have started to explore alternative approximation strategies to fit GLMM. For example, in bisulfite sequencing studies (Weissbrod, et al., 2017), the Mixed model Association via a Laplace ApproXimation algorithm (MALAX) relies on a Laplace approximation to improve computational speed. However, the computational improvement of MALAX over MACAU is relatively marginal (approximately two-folds). In non-genomics sequencing settings, a score test based approximate algorithm has also been recently developed to apply GLMM to analyze large-scale GWASs (Chen, et al., 2016). However, score test based strategy is not well-suited for genomic study setting where the null model varies for every genomic unit tested (e.g. gene or CpG site). Therefore, scaling up GLMM to thousands of individuals remains a challenging task.

In terms of accuracy, existing algorithms to fit GLMM rely on different approximations and these different approximations may work well in different settings. For example, in the field of biostatistics, it has been shown that while some GLMM algorithms may produce accurate *p*-values for differential analysis tasks in small studies, other GLMM algorithms rely on asymptotic properties of the likelihood and can only produce accurate p-values when sample size is relatively large (Breslow and Lin, 1995; Browne and Draper, 2006; Fong, et al., 2010; Jang and Lim, 2009; Lin and Breslow, 1996; Rodriguez and Goldman, 2001). Therefore, exploring the behavior of different GLMM algorithms in different settings will be informative for practitioners. In addition, as we will show below, existing GLMM algorithms in genomic sequencing studies cannot yet provide accurate heritability estimates.

Here, we develop a new method and a software tool to enable scalable and accurate inference with GLMM for large-scale RNAseq and BSseq studies. We also perform extensive simulations to comprehensively evaluate our method together with several other existing methods in various simulation settings to give out recommendations on GLMM based differential analysis and heritability estimation for practitioners. Our newly developed method is based on the penalized quasi-likelihood (PQL) approximation algorithm (Breslow and Clayton, 1993), applies to GLMM with two or more variance components, and with an efficient implementation, is capable of utilizing the parallel computing environment readily available in modern desktop computers. With the multiple-thread computing capability, our method can improve computation time for GLMM analysis of large-scale genomic sequencing data by at least an order of magnitude, making GLMM based differential analysis and heritability estimation applicable to hundreds or thousands of individuals. Importantly, as we will show below, our method is currently the only available method that can produce unbiased heritability estimates for sequencing count data. We refer to our method as the Penalized Quasi-Likelihood for sequencing count data (PQLseq). With extensive simulations and comparison with LMM or other existing GLMM methods, we illustrate both the advantage and limitation of our method. Finally, we apply our method to analyze a large-scale RNAseq study in the Hutterites.

## Materials and Methods

### PQLseq: Models and Algorithm

PQLseq fits two forms of GLMM that include the Poisson mixed model (PMM) for modeling RNAseq data and the binomial mixed model (BMM) for modeling BSseq data. These two different types of sequencing data have different data structures. Specifically, RNAseq studies collect one read count for each gene as a measurement of its expression level. In contrast, BSseq studies collect two read counts for each CpG site – one methylated count and one total count – as a measurement of the methylation level at the CpG site. The ratio between these two counts in the BSseq data represents approximately the methylation proportion of the given CpG site. Therefore, we use two different types of GLMM to model RNAseq and BSseq data. For both data types, we examine one genomic unit (i.e. gene or CpG site) at a time.

For a given gene in an RNAseq study, we consider the following PMM

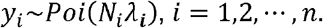

where *n* is the number of individuals; *y_i_* is the number of reads mapped to the particular gene for the *i*’th individual; *N_i_* is the total read counts for the individual (a.k.a read depth or coverage); and *λ_i_* is an unknown Poisson rate parameter that represents the underlying gene expression level for the individual.

For a given CpG site in a BSseq study, we instead consider the following BMM

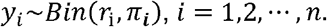

where *r_i_* is the total read count for *i*’th individual; *y_i_* is the methylated read count for that individual, constrained to be an integer value less than or equal to *r_i_*; and *π_i_* is an unknown parameter that represents the underlying proportion of methylated reads for the individual at the site.

For either model, we transform the unknown parameters into a latent variable *z_i_*: *z_i_* = log (*λ_i_*) in PMM and *z_i_* = logit (*π_i_*) in BMM. We then model the latent variable *z_i_* as a linear combination of several parameters,

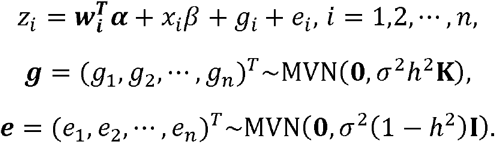

where *w_i_* is a c-vector of covariates including the intercept; *α* is a c-vector of corresponding coefficients; *x_i_* represents the predictor variable of interest (e.g. experimental perturbation, sex, disease status, or genotype); *β* is its ***g*** coefficient; is an *n*-vector of genetic effects; **e** is an *n* -vector of environmental effects; **K** is an *n* by *n* relatedness matrix that models the covariance among individuals due to either individual relatedness or population structure; **I** is an *n* by *n* identity matrix that models independent environmental variation; *σ^2^h*^2^ is the genetic variance component; *σ^2^(1 – h*^2^) is the environmental variance component; and MVN denotes the multivariate normal distribution. When **K** is standardized to have *tr*(**K**)/*n* = 1,*h^2^* ∈ [0,1] has the usual interpretation of heritability (Zhou, et al., 2013; Zhou and Stephens, 2012), where the *tr*(.) denotes the trace of a matrix.

In the above BMM and PMM, we are interested in testing the null hypothesis ***H*_0_**:*β* = 0 and/or estimating the heritability parameter *h^2^*. Both tasks require the development of computational algorithms to fit GLMM. Unfortunately, fitting GLMM is notoriously difficult, as the GLMM likelihood consists of an n-dimensional integral that cannot be solved analytically. To overcome the high dimensional integral and enable scalable estimation and inference with GLMM, we develop an approximate fitting method based on the penalized quasi-likelihood (PQL) approach (Breslow and Clayton, 1993) that is also recently applied to GWAS settings (Chen, et al., 2016). The detailed algorithm is provided in the Supplementary Text. Briefly, our method employs an iterative numerical optimization procedure. In each iteration, we introduce a set of pseudo-data 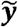 to replace the originally observed data ***y***. The pseudo-data 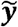 is obtained based on a second order Taylor expansion using the conditional distribution *P(y|g,e)* using the first and second order moments *E(**y**|**g**,**e**)* and *V(**y**|**g**,**e**)*, both evaluated at the current estimates of the fixed coefficients *α* and *ß* as well as the random effects ***g*** and ***e***. With the pseudo-data, the complex GLMM likelihood function for the original data ***y*** is replaced by a much simpler LMM likelihood function for the pseudo-data 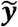 thereby alleviating much of the computational burden associated with GLMM. With the pseudo-data 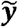 we can perform inference and update parameters using the standard average information (AI) algorithm for LMMs (Chen, et al., 2016; Gilmour, et al., 1995; Yang, et al., 2011). By iterating between the approximation step of obtaining the pseudo-data 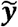 and the inference step of updating the parameter estimates via the AI algorithm, the PQL approach allows us to perform inference in a computationally efficient fashion. To improve computational speed further, we also take advantage of the parallel computing environment readily available in modern desktop computers nowadays and implement our method with multiple-thread computing capability using Rcpp. We refer to our method as the Penalized Quasi-Likelihood for sequencing count data (PQLseq), which is freely available as an R package at www.xzlab.org/software.html and https://cran.r-project.org/web/packages/PQLseq/index.html.

### Simulations

We performed simulations to compare different methods. To make simulations as realistic as possible, we simulated either RNAseq data or BSseq data based on parameters inferred from two published data sets that include a RNAseq data set (Tung, et al., 2015) and a BSseq data set (Lea, et al., 2015). In the simulations, we varied the sample size (*n*) (*n* = 50, 100, 200, 300, or 500). To construct a relatedness matrix ***Κ*** in each of these sample simulations, we first obtained a real relatedness matrix from the published data (Lea, et al., 2015). We then constructed the relatedness matrix ***Κ*** by filling in its off-diagonal elements with randomly drawn off-diagonal elements from the real relatedness matrix following (Lea, et al., 2015). In cases where the resulting ***Κ*** was not positive definite, we used the *nearPD* function in R to find the closest positive definite matrix as the final ***K***. Besides *n* and ***K***, we also simulated a continuous predictor variable ***x*** from a standard normal distribution, and normalized the predictor *x* to have a zero mean and unit variance.

For RNAseq based simulations, in each simulation replicate, we simulated the total read count *N_i_* for each individual from a discrete uniform distribution with a minimum (=1,770,083) and a maximum (=9,675,989) total read count (i.e., summation of read counts across all genes) equal to the minimum and maximum total read counts in the published RNAseq data (Tung, et al., 2015). We simulated 10,000 gene expression values and considered two general simulation settings. In the null settings, we simulated 10,000 non-differentially expressed (non-DE) genes to examine the gene expression heritability estimation accuracy and type I error control. In the alternative settings we simulated 1,000 DE genes and 9,000 non-DE genes to examine power. These non-DE or DE genes are simulated using the following procedure. Specifically, for each gene in turn, we simulated the genetic random effects ***g*** from a multivariate normal distribution with covariance ***Κ***. We simulated the environmental random effects ***e*** based on independent normal distributions. We then scaled the two sets of random effects to ensure a fixed value of heritability 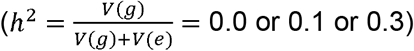 and a fixed value of over-dispersion variance (σ^2^ =*V(g) + V(e)* = 0.1, 0.25 or 0.5) where the function *V*(·) denotes the sample variance. A heritability value of 0.1 and 0.3 correspond approximately to the median and upper 15% percentile of gene expression heritability estimates from the RNAseq data (Tung, et al., 2015). An over-dispersion variance value of 0.1, 0.25 and 0.5 correspond to approximately the lower quartile, median, and upper quartile of the over-dispersion variance inferred from the RNAseq data (Tung, et al., 2015). Afterwards, for non-DE genes, the genetic effects ***g***, environmental effects ***e***, and an intercept (µ) were then summed together to yield the latent variable log(λ) = μ + ***g*** + ***e***. Here, the intercept 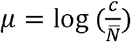 ensures an average gene count of *c* = 10, 50, or 100, where is the average total read count across individuals. For DE genes, we used log(λ) = μ + ***x***β + ***g*** + ***e*** to yield the latent variable, where 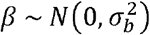 and 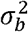 is set to ensure a fixed proportional of variance explained (PVE). That is, 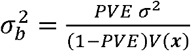 where PVE values were fixed to be 15%, 25%, or 35% to represent different effect sizes. Finally, we simulated the read counts based on a Poisson distribution with the Poisson rate being a product of the total read counts *N_i_* and the latent variable *λ_i_*; that is,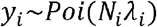 for the *i*’th individual. With the above procedure, we first simulated data under *n* = 100, *h^2^* = 0.1 and *σ*^2^ = 0.25 (and PVE = 0.25 for DE genes). We then varied one parameter at a time to generate different simulation scenarios. In each scenario, conditional on the sample size, total read counts etc., we performed 10 simulation replicates, each consisting of 10,000 genes.

For BSseq based simulations, in each simulation replicate, we simulated methylation values for 10,000 sites and considered two general simulation settings. In the null settings, we simulated 10,000 non-differentially methylated (non-DM) sites to examine the methylation level heritability estimation accuracy and type I error control. In the alternative settings we simulated 1,000 DM sites and 9,000 non-DM sites to examine power. These non-DM or DM sites are simulated using the following procedure. Specifically, for each site in turn, we simulated total read counts *r_i_* for each individual *i* from a negative binomial distribution *r_i_ ~ ΝΒ*(*μ,θ*) with *μ* = 18.80 and median *θ* = 2.49; the two parameter values correspond to the median estimates from the published BSseq data (Lea, et al., 2015). We then simulated the genetic random effects *g* and the environmental random effects *e* given a fixed heritability *h*^2^ (0.1 or 0.3) and a fixed value of over-dispersion variance (σ^2^ = 0.5, 1.2, or 2). Again, the over-dispersion variance values correspond to the lower quartile, median, and upper quartile of the over-dispersion variance inferred from the BSseq data (Lea, et al., 2015). For non-DM sites, the genetic effects, environmental effects, and an intercept (µ) were then summed together to yield the latent variable logit(π) =*μ + g +* e. Here, 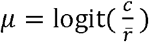 ensures an average number of methylated read counts being approximately c =5, 10 or 19, where r is the average total read count for the given site across individuals. For DM sites, we use logit(π) *= μ + xß + g + e* to yield the latent variable, where 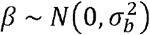 and 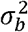 is set to ensure a fixed PVE. That is, 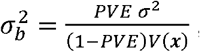, where PVE values were set to be 15%, 25%, or 35% to represent different effect sizes. Finally, we simulated the methylated read counts based on a binomial distribution with a rate parameter determined by the total read counts *r_i_* and the methylation proportion *π_i_*; that is, *y_i_*~*Bin*(*r_i_*,*π_i_*) for the *i*’th individual. With the above procedure, we first simulated data under *n* = 100, *h^2^ =* 0.1 and σ^2^ = 1.2 (and PVE = 0.15 or PVE = 0.25 for DM sites). We then varied one parameter at a time to generate different scenarios. In each scenario, conditional on the sample size, total read counts etc., we performed 10 simulation replicates, each consisting of 10,000 sites.

We compared four different methods (PQLseq, MACAU, GEMMA, and MALAX) in the BSseq based simulations, and compared three different methods (PQLseq, MACAU and GEMMA) in the RNAseq based simulations as MALAX is only applicable for BSseq data. For GEMMA, we normalized data following previous recommendations (Lea, et al., 2015; Sun, et al., 2017). Specifically, for RNAseq data, for each gene in turn, we divided the number of read counts mapped to the gene by the total read depth, and quantile transformed the normalized data to a standard normal distribution. For BSseq data, we used “M” value transformation following (Du, et al., 2010) by dividing the number of methylated reads by the number of unmethylated reads followed by a log2-transformation. The normalized data is 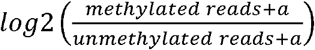, where *a* = 0.01 to avoid log transforming zero values.

### Real Data Application

The published RNAseq data was collected from lymphoblastoid cell lines (LCLs) of 431 individuals from the Hutterites population in South Dakota, which is an isolated founder population (Cusanovich, et al., 2016). Libraries were created using the TruSeq Library Kit and samples were sequenced an Illumina HiSeq 2000 (50bp single end reads) in indexed pools of 12. Reads were trimmed for adaptors using Cutadapt (reads less than 5 bp discarded) then remapped to hg19 using bowtie indexed with gencode version 19 gene annotations (Langmead, et al., 2009; Martin, 2011). To remove mapping bias, autosomal reads were processed through WASP (van de Geijn, et al., 2015). Gene counts were quantified using HTSeq-count (Anders, et al., 2015) and verifyBamID was used to identify sample swaps (Jun, et al., 2012). Following these mapping and quality control steps, we obtained expression count measurements for 23,367 genes. We kept genes that have read counts greater than five in at least two individuals to focus on a final set of 17,312 genes. We also used the Hutterites pedigree information to compute the kinship coefficients between pairs of individuals and used them as the ***K*** if matrix in the model. We then fitted a PMM for each gene in turn without any covariates to estimate gene expression heritability. For DE analysis, we used sex as the predictor variable (i.e. male vs female) to identify sex-associated genes. To compare performance among different methods for DE analysis, we permuted phenotype sex 20 times to obtain a null distribution. We used the null distribution to estimate the false discovery rate (FDR). Finally, to explore the influence of batch effects for heritability estimates in PQLseq, we extracted the top principal components (PCs) from the gene expression matrix and treated them as covariates in the model. We considered including a different number of top gene expression PCs that range from 2 to 200.

## Results

We provide a brief overview of the PQLseq method in the Materials and Methods section, with algorithmic details available in the Supplementary Text. Briefly, PQLseq fits a Poisson mixed model (PMM) for modeling RNAseq data and a binomial mixed model (BMM) for modeling BSseq data. In both data types, PQLseq examines one genomic unit (i.e. a gene or a CpG site) at a time, produces an estimated heritability 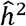, and in the case of differential analysis, also computes a *p*-value for testing the genomic unit association with a predictor variable of interest, where the predictor variable can be either continuous or discrete.

We performed a series of simulations to compare the performance of PQLseq with three other commonly used methods: (1) a linear mixed model implemented in GEMMA (Zhou, et al., 2013; Zhou and Stephens, 2012); (2) a GLMM model fitted using an MCMC based p-value computation algorithm implemented in MACAU (Lea, et al., 2015; Sun, et al., 2017); and in the case of BSseq based simulations, we also compared with (3) a binomial mixed model fitted using a Laplace approximation algorithm implemented in MALAX that is specifically designed for analyzing BSseq data. We did not compare with other commonly used DM or DE methods because (1) our main focus here is on examining the performance of different GLMM algorithms and LMM methods, (2) extensive simulations comparing other DM or DE methods with GLMM have been carried out elsewhere (Lea, et al., 2015; Sun, et al., 2017). Here, we simulated RNAseq data or BSseq data based on parameters inferred from published data sets (Lea, et al., 2015; Tung, et al., 2015) (simulation details in the Materials and Methods). In particular, in each setting, we simulated methylation levels for 10,000 sites or gene expression levels for 10,000 genes. In all these cases, the simulated gene expression levels or methylation levels are influenced by both independent environmental effects and correlated genetic effects, where the genetic effects are simulated based on a kinship matrix with either zero (*h*^2^ = 0.0), moderate (*h*^2^ = 0.1), or high (*h*^2^ = 0.3) heritability values. A heritability value of 0.1 corresponds approximately to the median heritability estimate in our real data analysis (see below), while a heritability value of 0.3 corresponds approximately to the 85th percentile of the expression heritability estimated in the real data (Lea, et al., 2015; Tung, et al., 2015).

### PQLseq produces approximately unbiased heritability estimates for sequencing count data

Our first set of simulations was performed to evaluate the effectiveness of PQLseq in terms of heritability estimation. To do so, we simulated BSseq data or RNAseq data with a fixed heritability value that equals to either 0.1 or 0.3. We considered sample sizes ranging from *n =* 50 to *n =* 500. We varied mean observed read count values (μ) and over-dispersion variance values (σ^2^) from low, moderate, to high, in order to examine how these parameters impact heritability estimation accuracy. Heritability estimates from different methods for different sample sizes in the BSseq based simulations are shown in Figures 1A and 1B. Heritability estimates for RNAseq based simulations are shown in Figures 1C and 1D. Heritability estimates from different methods for increasing μ are shown in Figure S1 and estimates for increasing σ^2^ are shown in Figure S2.

**Figure 1.**
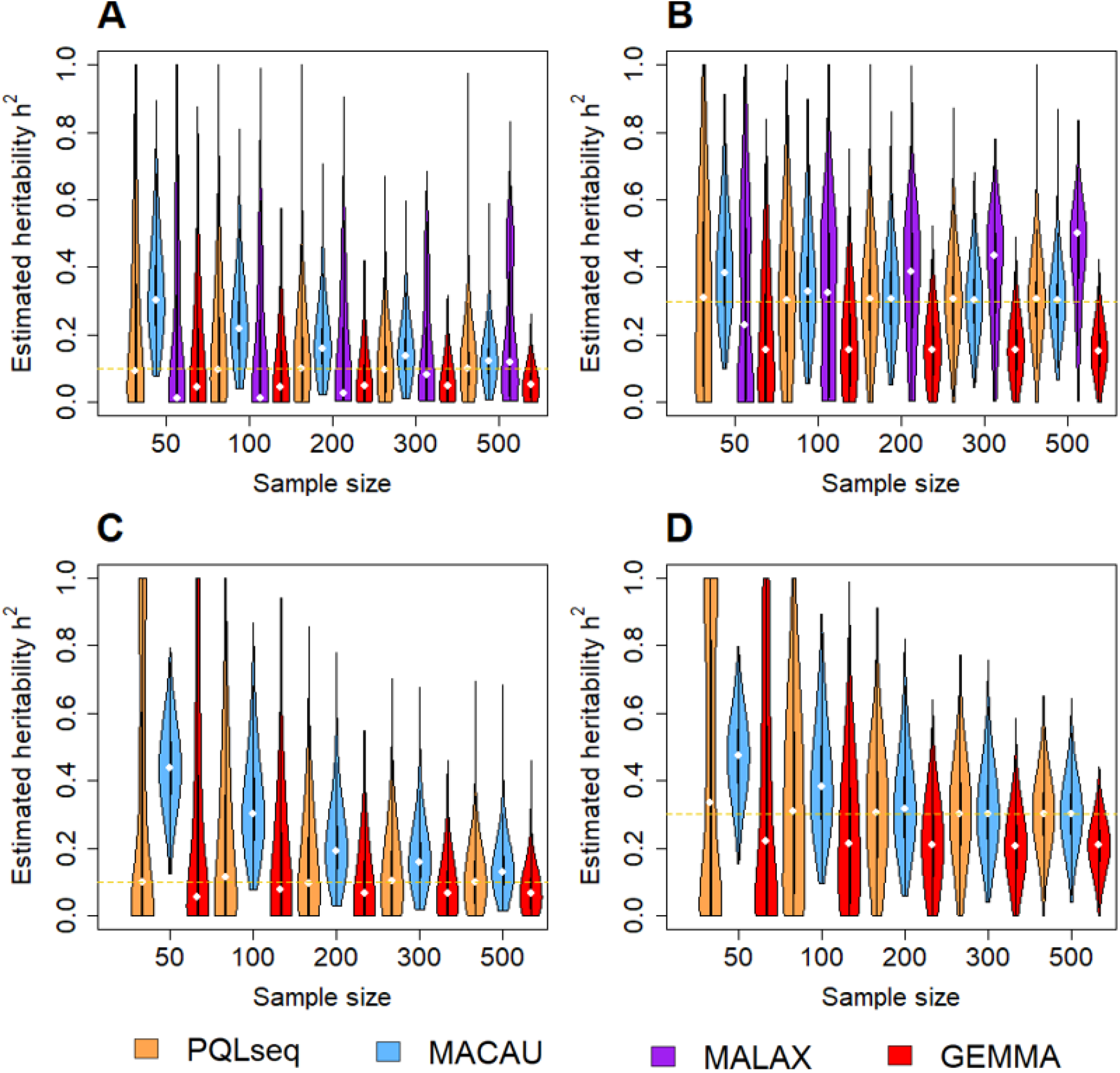
PQLseq produces unbiased heritability estimates across a range of sample sizes in BSseq and RNAseq based simulations. Violin plots display heritability estimates obtained from PQLseq (orange), MACAU (blue), MALAX (purple), and GEMMA (red). The first two panels show heritability estimates from PQLseq, MACAU, MALAX, and GEMMA in BSseq based simulation with parameters = 1.2, = 19, and with = 0.1 **(A)** or = 0.3 **(B).** The second two panels show heritability estimates from PQLseq, MACAU, and GEMMA in RNAseq based simulation with parameters = 0.25, and with = 0.1 **(C)** or = 0.3 **(D).** The horizontal orange dashed line represents the true heritability.

Across all simulation settings, PQLseq is the only method that produces approximately unbiased heritability estimates. In contrast, LMM implemented in GEMMA consistently produces downward biased estimates across different sample sizes, and more so for high heritability values (i.e., 0.3) than for low heritability values (i.e., 0.1). The downward bias of LMM presumably stems from the fact that LMM fails to model the count data generating process and inappropriately attributes the count generating noise to environmental errors. Increasing sample size (Figure 1) or over-dispersion variance (Figure S2) does not alleviate the downward bias of GEMMA. However, increasing the mean observed read counts alleviates the LMM estimation bias in RNAseq data (Figure S1), presumably because the normal approximation in LMM becomes appropriate with high read counts. On the other hand, MACAU produces consistently upward biased estimates across sample sizes, and more so for low heritability values (i.e. 0.1) than for high heritability values (i.e. 0.3). Increasing the observed read counts (Figure S1) or over-dispersion variance (Figure S2) does not alleviate the upward bias of MACAU. The upward bias in MACAU presumably stems from its inaccurate latent variable approximation algorithm in small samples. Indeed, with increasing sample size (e.g. *n* > 200), the heritability estimates from MACAU become approximately unbiased. For BSseq based simulations, MALAX also produces biased heritability estimates. The heritability estimates from MALAX is highly dependent on sample size: they are downward biased in small samples and becomes upward biased in large samples (Figure 1). Increasing the observed read counts (Figure S1) or over-dispersion variance (Figure S2) does not alleviate the heritability estimation bias of MALAX. Certainly, due to well-known drawbacks of PQL (Lin, 2007; Lin and Breslow, 1996), the variance component estimates in terms of ‼^2^ can display downward bias in extreme small sample sizes (e.g. *n* = 10, 20; Figure S3), which further affects the estimation of heritability *h*^2^. However, estimation of is impractical in these extreme small samples anyway due to the large standard errors there. Overall, our results suggest that PQLseq is the only method currently available that can produce approximately unbiased heritability estimates for sequencing count data with reasonable sample size.

Next, in addition to heritability estimation, we also examined the use of PQLseq for SNP heritability estimation. In particular, we examined how different genetic architectures of gene expression might influence SNP heritability estimation results. To do so, we obtained genotype data from *n* = 465 individuals with European ancestry from the GEUVADIS study (Lappalainen, et al., 2013) as processed in (Zeng and Zhou, 2017). We extracted 810 SNPs within +/-10kb of a median-sized gene (LIN9) from the data and used these real genotypes to simulate gene expression phenotypes. In the simulations, we varied the proportion of causal SNPs from 2%, 10% to 100% to capture a wide range of sparse to polygenic genetic architectures. In each setting, we simulated causal SNP effects each from a normal distribution and summed their effects to form the genetic random effects term. We also simulated the residual errors from a normal distribution to form the environmental random effects term. We scaled both terms so that the causal SNPs in total account for a heritability of *h*^2^ = 0.1 or *h*^2^ = 0.3. Afterwards, we simulated count data based on the same parameter settings and procedure explained in the RNAseq based simulations above. In addition to the above GLMM approaches, we also applied the Bayesian sparse linear mixed model (BSLMM) (Zhou, et al., 2013), which is commonly used for SNP heritability estimation in various genetic architectures and which models normalized data. The results are shown in Figure S4 and are largely consistent with the comparative results on heritability estimation described in the above paragraph. In particular, GEMMA produces slightly downward biased estimates; BSLMM also requires normalized data and occasionally produces downward biased estimates (when the causal SNP proportion is 10% and *h*^2^ = 0.3); MACAU produces slightly upward biased estimates; while PQLseq performs reasonably well and yields approximately unbiased estimates in various polygenic settings.

### PQLseq provides effective control of type I error for differential analysis in large samples

Our second set of simulations was performed to evaluate the effectiveness of PQLseq in controlling for type I error for differential analysis under sample non-independence. Sample non-independence is a common phenomenon in sequencing studies and can be caused by individual relatedness, population structure, or hidden confounding factors (Dubin, et al., 2015; Scott, et al., 2016; Tung, et al., 2015). Failing to properly control for sample non-independence can lead to inflated type I errors (Lea, et al., 2015; Sun, et al., 2017). To examine type I error control of different methods, we performed null simulations and simulated outcome variables in terms of methylation levels or gene expression levels that are independent of the predictor variable of interest ***x*** However, these outcome values are correlated among individuals/samples, with correlation determined by a heritability value of either 0.1 or 0.3. We considered sample sizes ranging from *n* = 50 to *n* = 500 We also varied mean observed read count values (µ) and over-dispersion variance values (⃃^2^) from low, moderate, to high, in order to examine how these parameters impact type I error control. In these simulations, we examined one genomic unit (i.e. site or gene) at a time and computed *p*-values using different methods.

We first calculated the genomic control factors based on the p-values for each method at a time and display them across different sample sizes in Figure 2A (for BSseq based simulations) and Figure 2D (for RNAseq based simulations). Corresponding genomic control factors for increasing *μ* are shown in Figures S5A and S5D while the results for increasing *σ^2^* are shown in Figures S6A and S6D. Overall, the genomic control factors from PQLseq and GEMMA are closer to the expected value of one compared with the other two methods (MACAU and MALAX) across the different sample sizes. In contrast, the p-values from MACAU are slightly conservative in small samples with genomic control factors lying below one. The conservativeness of MACAU are consistent with previous studies (Lea, et al., 2015; Sun, et al., 2017) and presumably stems from inaccurate asymptotic approximation in small sample sizes. Indeed, the genomic control factor from MACAU quickly approaches one with increasingly large sample sizes. On the other hand, in BSseq based simulations, MALAX produces slightly anti-conservative p-values with genomic control factors close to 1.1 in small to moderate samples (*n* = 50 ~ 200; Figure 2), or in cases where the observed read counts (Figure S5) or the over-dispersion variance (Figure S6) is low. The genomic control factor from MALAX approaches one when *n* = 300 and is below one when *n* = 500.

**Figure 2.**
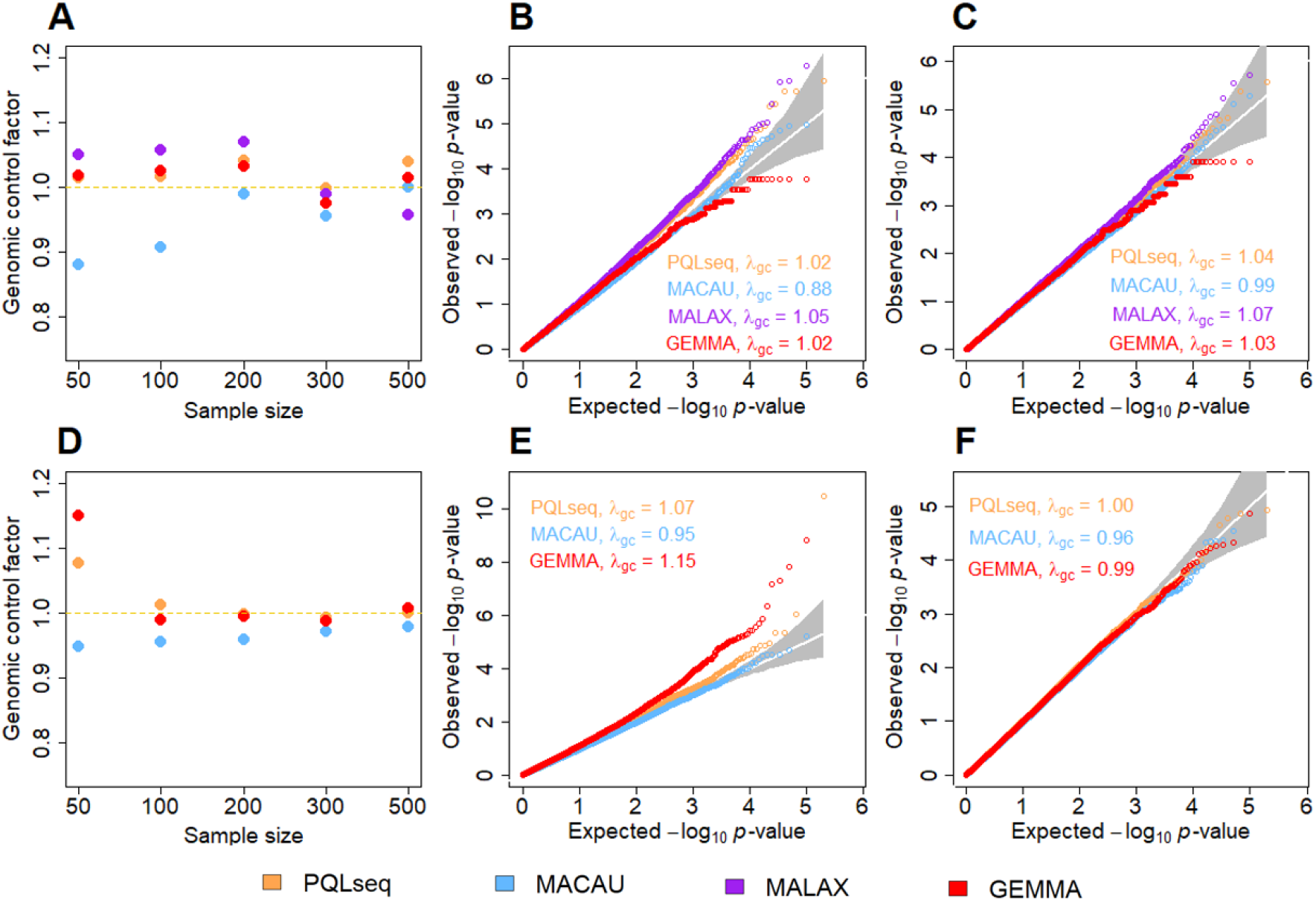
PQLseq produces calibrated *p*-values in BSseq and RNAseq based null simulations when sample size is large. Genomic control factors from PQLseq (orange), MACAU (blue), MALAX (purple) and GEMMA (red) across a range of sample sizes under the null are shown for BSseq based simulations **(A)** or RNAseq based simulations **(D).** Parameters used include = 1.2 and = 19 for BSseq based simulations and = 0.25 and = 10 for RNAseq simulations. QQ-plots further compare the expected and observed *p* -value distributions generated from different methods under the null aggregated from 10 simulation replicates for = 50 **(B)** and = 200 **(C)** in BSseq based simulations, and for = 50 **(E)** and = 200 **(F)** in RNAseq based simulations. is the genomic control factor.

In addition to genomic control factors, we also display QQ-plots of -log10 *p*-values from these methods in small samples (*n* = 50) in Figures 2B (for BSseq based simulations) and 2E (for RNAseq based simulations), and display QQ-plots of -log10 *p*-values from these methods in large samples (n = 200) in Figures 2C and 2F. The QQ-plots for small and large *μ* are shown in Figures S6B/S6E and S7C/S7F, respectively. The QQ-plots for small and large σ^2^ are shown in Figures S7B/S7E and S7C/S7F, respectively. Overall, GEMMA produces calibrated type I error control across most settings, which is consistent with previous studies (Lea, et al., 2015; Sun, et al., 2017). However, the p-values from GEMMA are slightly more significant than expected under the null in the RNAseq based simulations when the sample size is small (n = 50). While the genomic control factors from PQLseq is close to one across a range of sample sizes, we did notice in QQ-plots that the type I error rates of PQLseq are slightly anti-conservative in small samples with small deviation from the diagonal line for small p-values. For example, when η = 50, the type I error from PQLseq is 1.9×10^−3^ and 2.6×10^−4^ at a size of 1×10^−3^ and 1×10^−4^, respectively. The small inflation of *p*-value from PQLseq in small samples is presumably due to PQL’s inability to account for estimation uncertainty in the variance component parameters there, which is a known drawback of PQL (Breslow and Lin, 1995; Browne and Draper, 2006; Fong, et al., 2010; Goldstein and Rasbash, 1996; Jang and Lim, 2009; Lin and Breslow, 1996; Rodriguez and Goldman, 2001). However, the p-value inflation from PQLseq is no longer observed in large samples *(n* ≥ 100), regardless of the observed read counts (Figure S5) or over-dispersion variance (Figure S7). Consistent with the genomic factor inflation, MALAX also produces anti-conservative p-values in small samples, more so than PQLseq. For example, when η = 50, the type I error from MALAX is 2.3×10^−3^ and 3.9×10^−4^ at a size of 1×10^−3^ and 1×10^−4^, respectively. In contrast, MACAU produces conservative p-values which lines below the diagonal line in small samples. The p-values from MACAU become calibrated when sample size is large *(n* ≥ 100), regardless of the observed read counts (Figure S5) or over-dispersion variance (Figure S6). In addition, we also notice that the p-values computed from MALAX have a strong enrichment near 1, more so with increasing sample sizes (*n* > 200; Figure S7). Finally, we found that the p-values from PQLseq are highly correlated with that from MACAU across a range of sample sizes (*r*^2^ varies from 0.96 to 0.99; Figure S8), with increasingly large correlation for increasingly large sample size.

Overall, our null simulation results show that different GLMM methods can be either conservative (MACAU) or anti-conservative (PQLseq and MALAX) in small samples. However, all methods can produce calibrate type I error control in reasonably sized samples (PQLseq and MALAX for *n* ≥ 100; MALAX for *n* ≥ 300). The effectiveness of PQLseq in controlling for type I error in moderate to large samples suggest that PQLseq is particularly well suited for differential analysis in large sequencing data.

### PQLseq exhibits similar power for differential analysis as MACAU

Our final set of simulations were performed to compare the power of different methods in differential analysis. To do so, we simulated 10,000 sites or genes among which 1,000 of them are DM sites or DE genes. For DM sites or DE genes, we varied the effect sizes of predictor values so that the predictor variable explains a certain proportion of phenotypic variance (PVE = 15%, 25% or 35%). We examined the power of different methods to detect DM sites or DE genes based on a fixed false discovery rate (FDR). The power with respect to different sample sizes at an FDR of 10% for different heritability values are shown in Figure 3 (A-C for BSseq based simulations; D-F for RNAseq based simulations). The corresponding results for an FDR of 5% are shown in Figure S9. The power with respect to different PVE values are shown in Figure S10, with respect to different observed read counts are shown in Figure S11, with respect to different over-dispersion variance values are shown in Figure S12.

**Figure 3.**
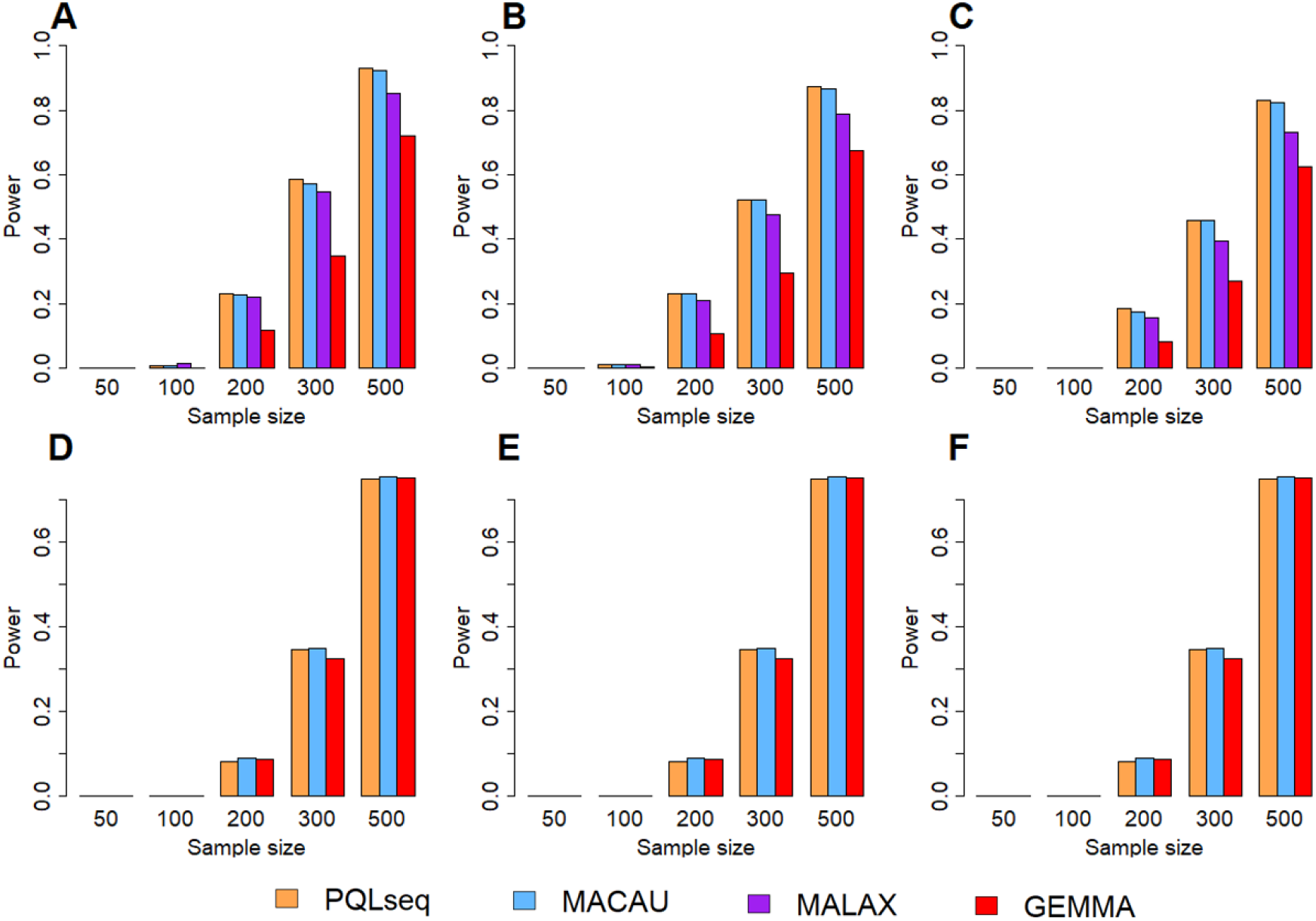
PQLseq exhibits similar power as MACAU in BSseq and RNAseq based power simulations across a range of sample sizes and heritability values. The power results are obtained for PQLseq (orange), MACAU (blue), MALAX (purple) and GEMMA (red) based on 10% FDR in both BSseq based simulations **(A, B, C)** and RNAseq based simulations **(D, E, F).** Results are shown under different heritability values: = 0 **(A** and **D),** = 0.1 **(B** and **E),** or = 0.3 **(C** and **F).** The other parameter settings in the simulations are = 19, PVE = 0.15 and = 1.2 for BSseq simulations; = 10, PVE = 0.25 and = 0.25 for RNAseq simulations.

In both BSseq and RNAseq based simulations, we found that all three GLMM methods (PQLseq, MACAU and MALAX) are more powerful than LMM method (GEMMA) across a range of simulation settings. The higher power of GLMM methods comes from their proper modeling of sequencing count data as demonstrated in previous studies (Lea, et al., 2015; Sun, et al., 2017). Among the different GLMM methods, we found that the performance of PQLseq, MACAU and MALAX are almost identical to each other when sample size is small (*n* ≤ 300), regardless of heritability values (Figure S9), PVE (Figure S10), read counts (Figure S11) and over-dispersion variance (Figure S12). The similarity in power between MACAU and MALAX in BSseq based simulations are consistent with the previous study (Weissbrod, et al., 2017). However, MACAU/PQLseq can be slightly more powerful than MALAX when sample size is large (*n ≥* 200; Figures 3A-C and S9). For example, when *η =* 200, *h^2^ = 0,PVE =* 0.15, at an FDR of 10%, we identified 230 DM sites with PQLseq, 226 with MACAU, but only 219 with MALAX. The power of MALAX decreases, however, with higher heritability. For example, when 0.3, *PVE =* 0.15, at an FDR of 10%, we identified 183 DM sites (whole significant sites) with PQLseq, 175 with MACAU, and 156 with MALAX. The reduced performance of MALAX in large samples as compared with other GLMM methods presumably is due to the unusual enrichment of MALAX p-values near one in large samples (Figure S7). The power comparison results also suggest that, despite the difference in type I error control, both PQLseq and MACAU rank genes or sites similarly well in terms of their differential expression or differential methylation evidence, thus producing similar power at a fixed FDR for differential analysis.

### PQLseq is computationally efficient

Finally, we emphasize that PQLseq is computationally efficient. For example, on a single CPU thread, PQLseq is 1.7 to 3 times faster than MACAU in both RNAseq and BSseq based simulations. In BSseq data, PQLseq is also comparable to MALAX. However, because PQLseq can take advantage of the multi-thread computing environment commonly available in modern computers, it can be an order of magnitude faster than MALAX and two orders or more of magnitude faster than MACAU (Figure 4). For example, it takes PQLseq, MACAU and MALAX 69.9, 182.2 and 46.6 hours, respectively, to analyze a data with 10,000 sites and *n* = 1,000 samples. However, with 10 threads, PQLseq can analyze the same sized data within 7.2 hours.

**Figure 4.**
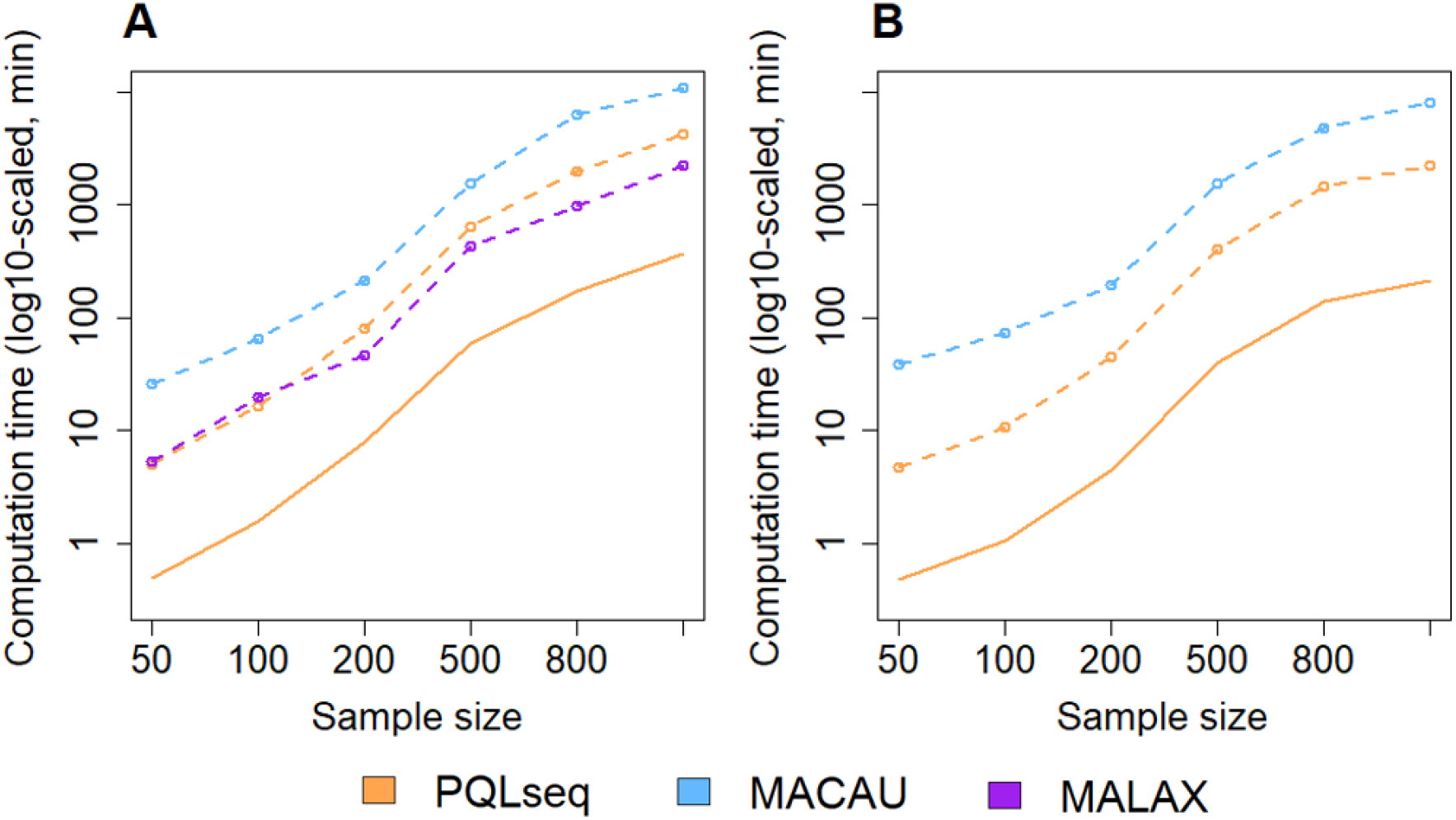
Computational time of different methods for analyzing data with different sample sizes. Plots show computational time, in minutes (log10-scaled), by MACAU (blue), MALAX (purple), PQLseq (orange, dashed line), or PQLseq with 10 threads (orange, solid line) across a range of sample sizes for analyzing 10,000 sites in BSseq based simulations **(A)** or 10,000 genes in RNAseq based simulations **(B).** Computation are carried out using Intel Xeon E5-2683 2.00 GHz processors.

### Analyzing the Hutterites RNA sequencing data

We applied PQLseq to analyze a published RNAseq data on 431 individuals from the Hutterites population in South Dakota, which is an isolated founder population (Cusanovich, et al., 2016). The Hutterites RNAseq data consists of gene expression measurements of 17,312 genes in lymphoblastoid cell lines (LCLs) from 431 individuals. These individuals are related. Specifically, 7,638 pairs of individuals in the data have a kinship coefficient exceeding 1/8 while 49,746 pairs have a kinship coefficient exceeding 1/16. To account for individual relatedness in the data, we applied LMM and GLMM based methods GEMMA, MACAU and PQLseq. We first used these methods to estimate expression heritability for all genes. Our heritability estimates are shown in Figure 5A. Specifically, the median heritability estimate across all genes is estimated to be 0.055 (mean estimate = 0.082) by GEMMA, 0.131 (mean = 0.162) by MACAU, and 0.071 (mean = 0.107) by PQLseq (Figure 5A). The order of these median estimates from different methods are consistent with the simulation results, with PQLseq estimate being higher than GEMMA and lower than MACAU. In addition, plotting individual expression heritability estimates from PQLseq against that from GEMMA or that from MACAU shows a similar pattern (Figure S13).

**Figure 5.**
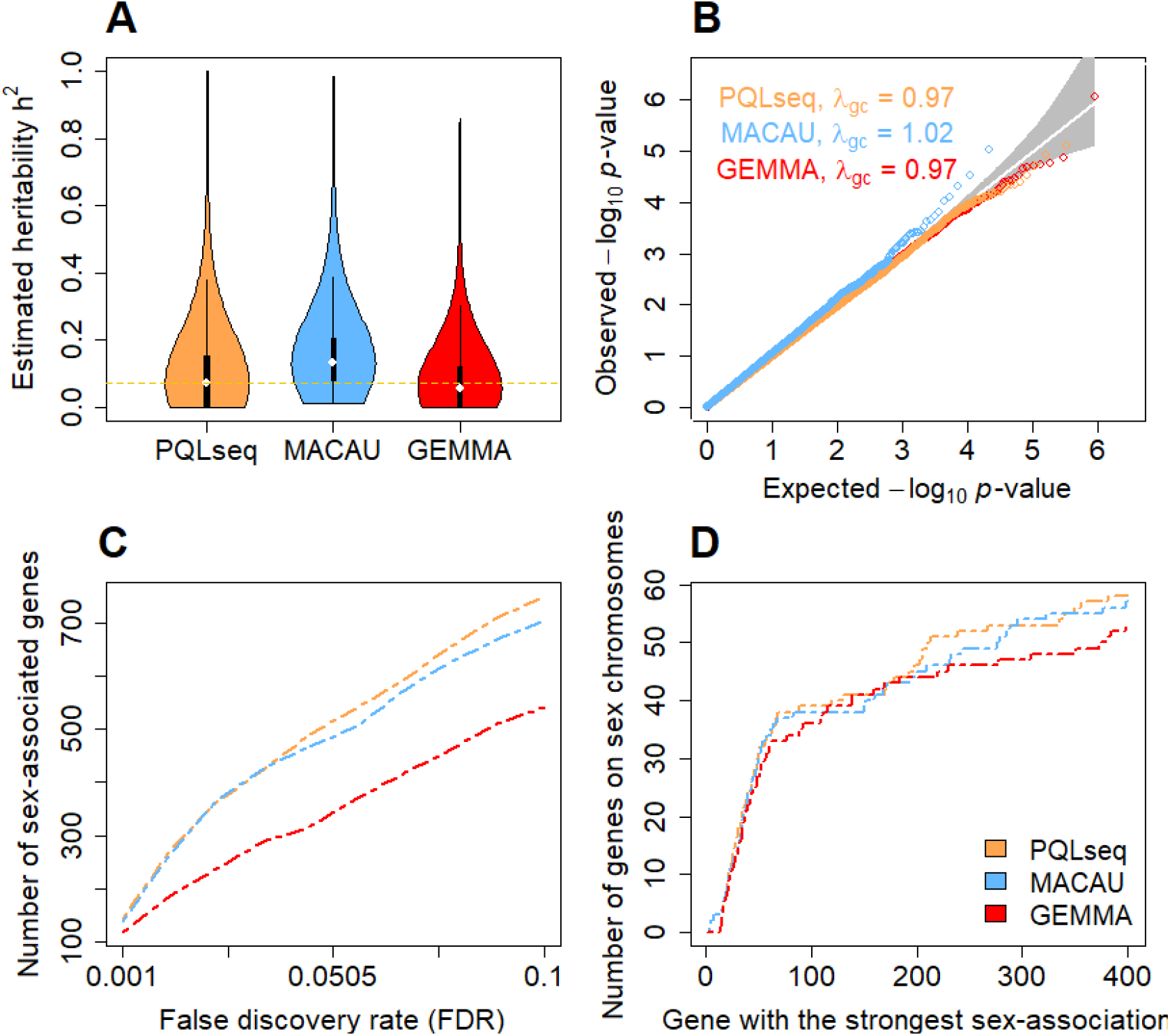
Gene expression heritability estimation and differential expression analysis in the Hutterites RNAseq data. Results are shown for PQLseq (orange), MACAU (blue) and GEMMA (red).**(A)** Violin plot shows gene expression heritability estimates from different methods. The median heritability estimate across genes is 0.071 for PQLseq, 0.131 for MACAU, and 0.055 for GEMMA. **(B)** QQ-plot comparing expected and observed *p*-values distributions under the permuted null for different methods. Results are aggregated across 20 permutations. *λ_gc_* is genomic control factor. **(C)** The number of genes that are associated with sex (*y*-axis) are plotted against false discovery rate (FDR) estimated based on permuted null (*x*-axis). **(D)** The number of genes that are on the sex chromosomes (*y*-axis) out of the genes have the strongest sex association (*x*-axis).

To explore the influence of batch effects on heritability estimates from PQLseq, we follow the original study (Cusanovich, et al., 2016) and extract the top principal components (PCs) from the gene expression matrix and treated them as covariates in the GLMM. Intuitively, removing batch effects would likely reduce measurement noise and subsequently improve heritability estimates. Indeed, we found that the *h* ^2^ estimates from PQLseq gradually increase with the addition of increasingly large number of PCs initially (Figure S15). The medium *h*^2^ estimate reach a plateau of 0.187 (mean = 0.249) near 160 PCs, which, however, is quite close to the medium *h*^2^ estimate of 0.165 (mean = 0.219) in the presence of 62 PCs – a number that maximizes number of expression quantitative trait (eQTL) discoveries in the original study (Cusanovich, et al., 2016). The *h*^2^ estimates from PQLseq gradually decrease after the plateau with the addition of more gene expression PCs, presumably because the later PCs do not necessary capture batch effects and may sometimes represent true biological/genetic effects.

Besides heritability estimation, we also performed DE analysis to detect genes whose expression level varies between genders (i.e. male vs female). The p-values for DE analysis are all well behaved (Figure S14). In order to compare methods based on a fixed FDR threshold, we also permuted the gender variable and performed DE analysis on the permuted data to construct an empirical null distribution for the p-values. In the permuted null data, all three methods produce calibrated p-values (Figure 5B), consistent with simulations. We then used the permuted null distribution of p-values to further estimate the empirical FDR at any p-value threshold and compared power of detecting sex-associated genes at a fixed FDR. The power comparison results are consistent with simulations and show that PQLseq and MACAU are more powerful than GEMMA. For example, at an empirical FDR of 10%, we identified 751 sex-associated genes with PQLseq, 706 sex-associated genes with MACAU, and 543 sex-associated genes with GEMMA (Figure 5C). For the two GLMM methods, the p-values from PQLseq are highly correlated with that from MACAU as expected (*r*^2^ = 0.91; Figure S16). We also verified that the top sex-associated genes identified by all three methods are enriched on sex chromosomes (Figure 5D), with PQLseq and MACAU showing slightly more enrichment than LMM, suggesting the detection of true associations (Lemos, et al., 2014; Vawter, et al., 2004; Zhou, et al., 2011). Finally, in terms of computation time, PQLseq finished the analysis in 1.2 hour with ten CPU threads while MACAU took 32 hours, suggesting that PQLseq is more computationally efficient than the previous GLMM method.

## Discussion

We have illustrated the benefits of using PQLseq to perform GLMM analysis on RNA sequencing and bisulfite sequencing data. We have shown that PQLseq is the only method currently available that can produce unbiased heritability estimates for sequencing count data. In addition, PQLseq is well suited for differential analysis in large sequencing studies, providing calibrated type I error control and more power than standard LMM methods. PQLseq is implemented as an R software package with parallel computing capacity, can accommodate both binary and continuous predictor variables, and can incorporate various biological or technical covariates as fixed effects. With simulations and real data applications, we have shown that PQLseq is a useful and efficient tool for analyzing genomic sequencing data sets that are becoming increasingly common and increasingly large.

In the paper, we have primarily focused on illustrating PQLseq for simple GLMMs with two-variance components: one component models sample non-independence due to the covariance matrix **K**, while the other component models independent over-dispersion. However, PQLseq can easily accommodate multiple variance components. Indeed, we have implemented PQLseq so that it can fit GLMMs with multiple variance components. GLMMs with multiple variance components can be particularly useful when there are multiple sources of variance that need to be accounted for (Weissbrod, et al., 2017). For example, one can use multiple variance components to account for population stratification, family relatedness as well as independent environmental variation. Alternatively, one can use multiple variance components to account for sample non-independence due to cell composition heterogeneity across samples, batch effects as well as independent noise. Exploring the use of GLMM with multiple variance components in various genomic sequencing studies is an interesting future direction.

In the present study, we have focused on analyzing large scale sequencing data with GLMM. Compared with small sample studies, sequencing studies with large sample sizes are better powered, more reproducible, and are thus becoming increasingly common in genomics (Ardlie, et al., 2015; Battle, et al., 2014; Consortium, 2015). For example, a recent comparative study makes explicit calls for moderate to large sample studies performed with at least 12 replicates per condition (i.e. *n* ≥ 24) (Schurch, et al., 2016). However, we recognize that many genomic sequencing studies are still carried out with a small number of samples (e.g. 3 replicates per condition). Estimating methylation or expression heritability in small samples is particularly challenging due to the high estimation uncertainty resulting from small samples. Indeed, even in our simulations with *n* = 50 samples, the heritability estimates are highly variable across simulation replicates (Figure 1). Therefore, we expect at least a couple hundred individuals are needed to yield reasonably accurate heritability estimates. For differential analysis, it is also well known that the power of all analysis methods can dramatically reduce with decreasing sample size, conditional on fixed values of other factors that influence power (e.g., effect size) (Lea, et al., 2015; Sun, et al., 2017). As a consequence, the advantage of GLMM over LMM may no longer be apparent in data with only three replicates per condition when the DE effect size is also kept small. Moreover, fitting GLMM in small data remains a challenging task: as we have shown in the present study, different GLMM algorithms can produce either conservative or anti-conservative p-values under the null in small samples. Therefore, exploring the use of other GLMM algorithms may help identify algorithms that are particularly well suited for small data. Studies have shown that the recently developed integrated nested Laplace approximation (INLA) algorithm can provide accurate parameter estimates in non-genomic settings (Holand, et al., 2013). While INLA is a Bayesian method, we can pair the INLA algorithm with the main idea of MACAU to rely on the difference of the posterior and the prior to enable frequentist estimation. By extracting the likelihood as the difference between the posterior and the prior, inference will no longer depend on the prior specification. Therefore, exploring the use of INLA or other GLMM algorithms may facilitate the application of GLMM to small data sets in the future.

Finally, as we have shown in the main text, while PQLseq is more computationally efficient than the other GLMM methods, its main computational gain over the other methods relies on multiple threads computation. It is certainly possible to implement the other algorithms to use parallel computation. In fact, because different GLMM algorithms can have different type I error control or sometimes different power for differential analysis in different settings, enabling parallel computation for other GLMM algorithms can provide more analytic options for practitioners. In the present study, we have only examined the computational scalability of PQLseq at a sample size up to *n* = 1,000, which is close to the largest genomic sequencing study performed thus far (n = 922) (Battle, et al., 2014). However, future studies will likely collect data with even larger samples. In addition, other genomic analysis such as molecular quantitative trait locus (QTL) mapping studies requires examining pairs of genes or sites with single nucleic polymorphisms (SNPs). Examining pairs of genes or sites with SNPs, even restricted to SNPs at the cis-regions of these genes or sites, will require a much larger number of tests than is required by heritability estimation or differential analysis. Applying GLMM to millions of tests for expression QTL (eQTL) or methylation QTL (meQTL) mapping, even with PQLseq and a relatively large computing cluster, is not a trivial task. Therefore, future algorithmic innovations are needed to scale up PQL or other algorithms to enable GLMM analysis both in larger data sets and for molecular QTL mapping studies.

## ACKNOWLEGMENTS

This study was supported by the National Institutes of Health (NIH) grants R01HG009124 and R01HL142023, and the National Science Foundation (NSF) grant DMS1712933. MC was supported by NIH grant R01GM126553. CO was supported by NIH grant R01HL085197. SS was supported by NIH grant R01HD088558 (PI Tung), Top International University Visiting Program for Outstanding Young scholars of Northwestern Polytechnical University, and the Fundamental Research Funds for the Central Universities (Grant NO. 3102017OQD098). JZ was supported by NIH grant U01HL137182 (PI Kang). We thank Dr. Noah Snyder-Mackler at Duke University for helping with parallel computation and thank William Wentworth-Sheilds for helping with the Hutterites data.

**Conflict of Interest:** none declared.

